# EphA7 Isoforms Differentially Regulate Cortical Dendrite Development

**DOI:** 10.1101/2020.03.27.011486

**Authors:** Carrie E. Leonard, Maryna Baydyuk, Marissa A. Stepler, Denver A. Burton, Maria J. Donoghue

## Abstract

The shape of a neuron reflects its cellular function and ultimately, how it operates in neural circuits. Dendrites receive and integrate incoming signals, including excitatory input onto dendritic spines, so understanding how dendritic development proceeds is fundamental for discerning neural function. Using loss- and gain-of-function paradigms, we previously demonstrated that EphA7 receptor signaling during cortical development impacts dendrites in two ways: restricting growth early and promoting spine formation later. Here, the molecular basis for this shift in EphA7 function is defined. Expression analyses reveal that both full-length (EphA7-FL) and truncated (EphA7-T1; lacking kinase domain) isoforms of EphA7 are expressed in the developing cortex, with peak expression of EphA7-FL overlapping with dendritic elaboration and highest levels of EphA7-T1 coinciding with spine formation. Overexpression studies in cultured neurons demonstrate that EphA7-FL inhibits both dendritic growth and spine formation, while EphA7-T1 increases spine density. Furthermore, signaling downstream of EphA7 varies during development; *in vivo* inhibition of kinase-dependent mTOR by rapamycin in EphA7 mutant neurons rescues the dendritic branching, but not the dendritic spine phenotypes. Finally, interaction and signaling modulation was examined. In cells in culture, direct interaction between EphA7-FL and EphA7-T1 is demonstrated which results in EphA7- T1-based modulation of EphA7-FL phosphorylation. *In vivo*, both isoforms are colocalized to cortical synapses and levels of phosphorylated EphA7-FL decrease as EphA7-T1 levels rise. Thus, the phenotypes of EphA7 during cortical dendrite development are explained by divergent functions of two variants of the receptor.

## Introduction

In the cerebral cortex, diverse neuronal populations are specifically arrayed and this organization underlies remarkable functional complexity (Rakic 1974; Angevine & Sidman 1961; Sidman & Rakic 1982; McConnell 1995). Indeed, distinct molecular and morphological profiles arise during development to determine functions of cortical projection neurons (Arlotta et al. 2005; Molyneaux et al. 2007). At embryonic timepoints, newly differentiated, bipolar-shaped neurons begin to migrate toward the cortical plate, with a leading process that draws the cell toward the pial surface and a trailing process that extends toward the ventricle (Anton et al. 1996; Noctor et al. 2004; Tsai & Gleeson 2005). This polarity sets the stage for the eventual maturation of the neuron, which is guided by a concert of intrinsic and extrinsic programs (reviewed in Whitford et al. 2002; Molyneaux et al. 2007). The trailing process becomes the axon and the leading process develops into the apical dendrite, which will retract or branch extensively, depending on the cell type. Then, additional basal dendrites branch from the soma (Miller 1981; Vercelli et al. 1992; Koester & O’Leary 1992; Polleux et al. 1998; Polleux et al. 2000; Guan & Rao 2003). As neurons elaborate and synaptic connections form during postnatal development, small, actin-rich protrusions from dendrites, called dendritic spines, serve as the postsynaptic site for excitatory synapses (Calabrese et al. 2006; Sala et al. 2008). Ultimately, dendritic morphology is cell type-specific, serving to integrate signals from specific afferents, and abnormal dendritic morphology has been linked to neurodevelopmental disorders (Häusser et al. 2000; Kaufmann & Moser 2000; Glausier & Lewis 2012; Phillips & Pozzo-Miller 2015; Ledda & Paratcha 2017). Despite recognition that dendritic form is critical to neuronal function, mechanisms guiding dendrite development remain obscure.

Previously, our research group demonstrated that the intercellular signaling molecule EphA7 plays a variety of roles in cortical neuronal development (Miller et al., 2006; Clifford et al. 2014). Membrane-bound Eph receptors engage ephrin ligands on an adjacent cell to affect cellular processes in one or both cells (Gale et al. 1996; Holland et al. 1996; Noren & Pasquale 2004). Analysis of neuronal maturation on patterned substrates revealed that EphA7 acts to restrict dendritic growth according to ligand distribution. Interestingly, Tsc1, a repressor of the common signaling intermediary, mammalian target of rapamycin (mTOR), is involved in EphA7 signaling (Clifford et al. 2014), consistent with the most well-characterized role of EphA7 as a repulsive molecule (Holmberg et al. 2000; Himanen et al. 2004; Torii & Levitt 2005; Miller et al. 2006; Lehigh et al. 2013). Yet, this previous data also demonstrated that EphA7 is capable of promoting dendritic spine formation, consistent with adhesive intercellular interactions in more mature EphA7- expressing neurons (Clifford et al. 2014). The mechanisms underlying diverse EphA7 functions from repulsive to adhesive signaling, however, were unclear.

The current study demonstrates that the opposing functions of EphA7 in dendritic growth and spine formation are mediated by two distinct isoforms produced by alternative splicing: a full-length EphA7 (EphA7-FL) that includes a kinase domain, and a truncated EphA7 (EphA7-T1) without enzyme activity (Valenzuela et al. 1995; Ciossek et al. 1995, Fig. 1A). Structurally, EphA7-FL and EphA7-T1 have identical extracellular regions, including ligand-binding and other protein interaction domains, as well as the transmembrane domain. Intracellularly, EphA7-FL contains a sterile alpha-motif, PDZ-binding domain, and a kinase domain, which, upon ligand-induced dimerization, is auto- phosphorylated, and initiates downstream signaling. In contrast, EphA7-T1 lacks intracellular signaling domains and, instead, has a unique 11 amino acid C-terminus (Ciossek et al. 1995, Fig. 1A). While previous studies demonstrated that EphA7-FL and EphA7-T1 are both expressed in developing brain, functional consequences of this expression were unknown (Ciossek et al. 1995; Mori et al. 1995; Valenzuela et al. 1995; Ciossek et al. 1999). A single study speculated that EphA7-T1 may contribute to adhesive cellular interactions during neural tube closure, demonstrating that ephrinA5-expressing non-neuronal cells mixed freely with EphA7-T1-expressing cells, but not EphA7-FL-expressing cells, and that the presence of EphA7-T1 decreased ligand-induced EphA7-FL phosphorylation (Holmberg et al. 2000). While the hypothesis that EphA7-T1 acts as a dominant negative against EphA7-FL was introduced, it remains untested in other systems and direct interaction between EphA7-FL and EphA7-T1 has yet to be proven.

**Figure 1.**
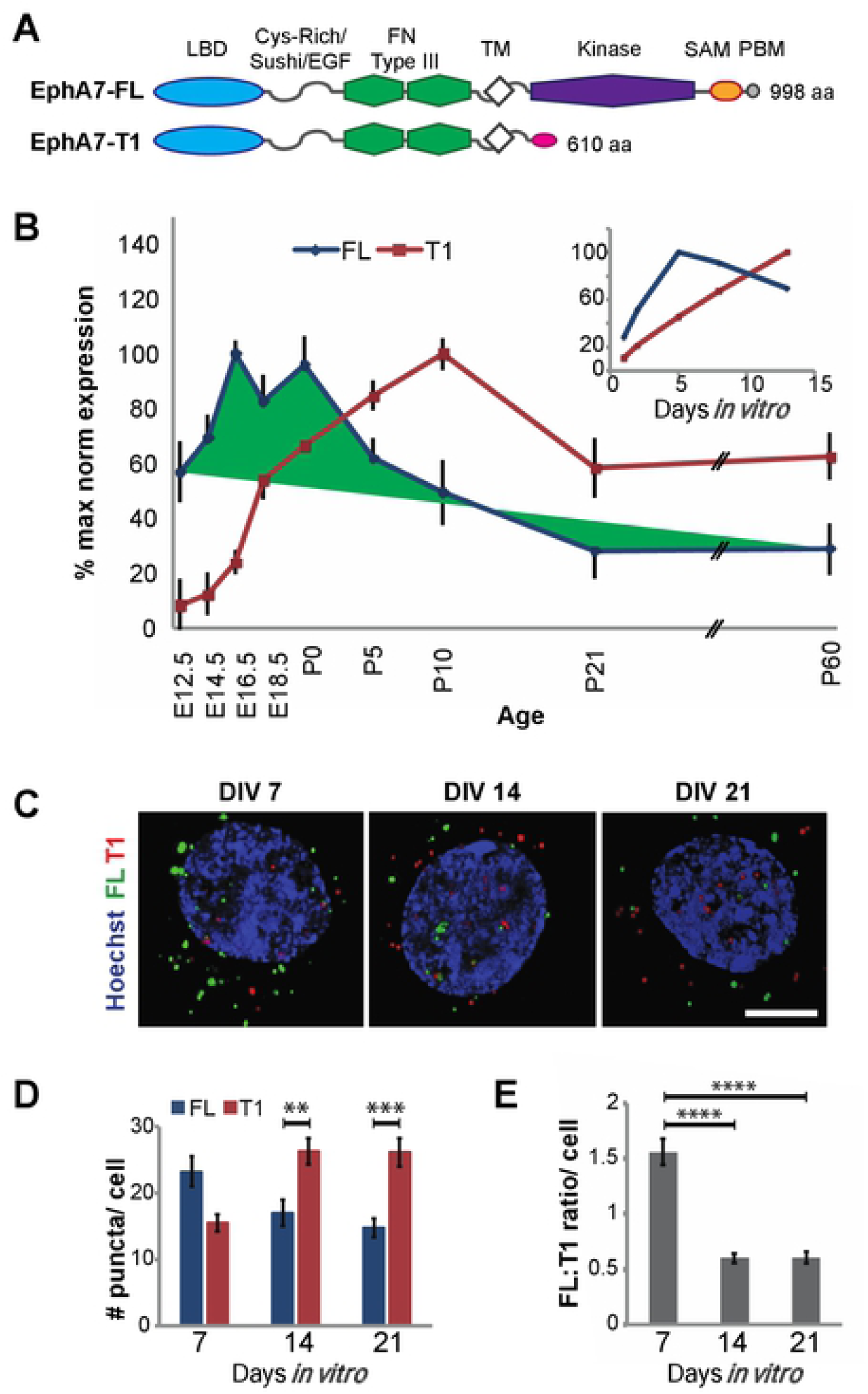
EphA7-FL and EphA7-T1 isoforms have dislinct and dynamic expression patterns during cortical development. A. Schematics of EphA7-FL and EphA7-T1 proteins. EphA7-FL (998 aa) and EphA7-TI (610 aa) have identical N-terminal regions, including ephrin ligand binding domain (LBD), cysteine-rich domain with Sushi and extracellular growth factor-like motifs (Cys-Rich/Sushi/EGF), fibronectin type-III repeats (FN Type III), and transmembrane regions (TM). However, EphA7-FL contains a kinase domain, sterile alpha motif(SAM), and PDZ-bindingdomain (PBD), while EphA7-T1 has a unique 11 aa C-tenninal end (pink). B. Levels of EphA7-FL (blue) and EphA7-T1 (red) transcripts during cortical development, as well as in cultured primary cortical neurons (inset), obtained with qRT-PCR and represented as percent maximum expression for each isoform, relative to U6 internal control. C. Fluoresce nt ISH corresponding to EphA7-FL (green) and EphA7-T1 (red) mRNA at single puncta resolution with the Hoechst-stained (blue) nuclei of primary cortical neurons at 7, 14, or 21 days *in vitro* (Scale bar, 5µm). D. Quantification of ISH puncta per cell revealed that the average number of EphA7-FL puncta per cell (blue bars) decreased from DIV7 to DIV21, while the average number of EphA7-T1 puncta (red bars) increased over time in culture. On average, there was a trend toward more EphA7-FL than EphA7-T1 puncta at DIV7 (p=0.06), and there were significantly more EphA7-T l than EphA7-FL puncta per cell at DIV14 and 21. E. The ratio of EphA7-FL to EphA7-T1 puncta per cell decreased from DIV7 to 14, with this ratio maintained at DIV 21. (** p<0.01, *** p<0.001, **** p<0.0001.

Here, unique expression patterns of EphA7 isoforms are characterized; expression of EphA7-FL or EphA7-T1 peak early during dendritic elaboration or later during spine formation, respectively. In addition, overexpressing EphA7-FL or EphA7-T1 produces discrete and opposing effects on dendrite maturation *in vitro*, with EphA7-FL-mediated dendrite restriction involving mTOR inhibition, but EphA7-T1-mediated spine formation being mTOR-independent. Finally, direct interaction between EphA7-FL and EphA7-T1 causes decreased phosphorylation of EphA7-FL, consistent with EphA7-T1 acting as a dominant negative, attenuating EphA7 repulsive signaling during dendritic spine formation. These results provide mechanistic details of EphA7 signaling during dendritic formation, and highlight the effects of developmentally controlled receptor isoforms in diversifying outcomes.

## Results

### EphA7 isoforms have distinct temporal expression in developing cortex

Cortical neurons in mutant mice lacking all *EphA7* isoforms (*EphA7*^*-/-*^) have longer, more complex dendrites and fewer excitatory synapses compared to cortical neurons in wild type (WT) mice (Clifford, et al. 2014). Previous studies have detected both EphA7-FL and EphA7-T1 (consisting of identical extracellular, but unique intracellular sequences) during normal cortical development, however detailed expression profiles have yet to be described (Valenzuela et al. 1995; Ciossek et al. 1995; Mori et al. 1995; Ciossek et al. 1999, Miller et al. 2006). To understand the developmental expression of EphA7-FL and EphA7-T1 in cortex, qRT-PCR was performed using isoform-specific primers. The EphA7-FL transcript was highly expressed during cortical development before birth, with peak expression between embryonic day (E) 16.5 and P0 and decreased in postnatal cortex (Fig.1B, blue). In contrast, EphA7-T1 transcript levels in the cortex were low throughout the embryonic period, increased during postnatal life with highest levels at P10 and then decreased, but stayed considerably higher than embryonic levels into adulthood (Fig.1B, red). A similar pattern of maturation-dependent isoform expression was observed in samples derived from cultures of primary cortical neurons (Fig.1B, inset).

To determine whether EphA7 isoforms are expressed in the same cells or in discrete populations, RNAscope *in situ* hybridization was used to detect mRNA puncta corresponding to each isoform. Probe specificity was confirmed by comparing fluorescence for each probe in WT versus *EphA7 KO* tissue (Supp. Fig.1A). EphA7- FL and EphA7-T1 transcripts were examined in individual neurons of primary cortical cultures at 7, 14, or 21 days *in vitro* (DIV, Fig. 1C). Immunofluorescent labeling of these cultures with NeuN (neuronal marker) or GFAP (glial marker) demonstrated that EphA7 transcripts were co-localized with NeuN and absent from cells expressing GFAP, indicating that EphA7 is primarily expressed in neurons at these stages (Supp. Fig.1B).

Quantification of puncta corresponding to EphA7-FL and EphA7-T1 transcripts demonstrated that early in cortical neuronal maturation, at DIV7, the average number of puncta/neuron for EphA7-FL trended higher than for EphA7-T1 (23 ±2 EphA7-FL vs 16 ±1 EphA7-T1, p=0.06, Fig.1C-E). The situation was reversed, however, as neurons matured; at both DIV14 and DIV21 the average number of puncta/neuron for EphA7-FL was lower than for EphA7-T1 (17±2 EphA7-FL vs 26 ±2 EphA7-T1 at DIV14, p<0.0001; 15 ±1 EphA7-FL vs 26 ±2 EphA7-T1 at DIV21, p<0.0001, Fig.1C-E). These analyses revealed that neurons positive for one EphA7 isoform also expressed the other, such that every cell positive for either EphA7-FL or EphA7-T1 mRNA was also positive for the other splice varient. The developmental shift in expression was also observed *in vivo* in motor cortex, where expression was high, with EphA7-FL predominantly expressed at P0 and EphA7-T1 preferentially expressed at P10 (Supp. Fig.1C-E). Taken together, these analyses of EphA7 expression reveal that EphA7-FL and EphA7-T1 are co-expressed within individual neurons, with higher proportions of EphA7-FL expression in immature neurons and more EphA7-T1 as neurons mature.

### Overexpression of EphA7-FL or EphA7-T1 *in vitro* results in divergent shifts in dendritic complexity and spine density

The discrete temporal expression of EphA7-FL and EphA7-T1 in maturing cortical neurons is consistent with the idea that the isoforms might differentially impact dendritic branching and spine maturation. To examine the functions of each isoform, epitope-tagged EphA7-FL (hemagglutinin for EphA7-FL; EphA7-FL-HA) and EphA7-T1 (myc for EphA7-T1; EphA7-T1-myc) expression constructs were created and their abilities to produce membrane-bound protein capable of binding ephrin-A5 ligand verified (Supp. Fig. 2). Then, primary cortical neurons were transfected at DIV7 (empty vector, EphA7-FL-HA or EphA7-T1-myc with actin-GFP to discern morphology). After harvesting neurons at DIV 21 and visualizing GFP, Sholl analysis was performed to quantify the complexity of dendritic arbors. In parallel, dendrites were examined to assess spine density. Compared to control-transfected neurons, dendrites of cells transfected with FL were less complex (Fig.2A,B) and had significantly lower dendritic spine density (control= 0.708 ± 0.03 spines/µm control; EphA7-FL-transfected= 0.476 ± 0.03 spines/µm, p<0.001, Fig.2C,D). These results indicate that elevation of EphA7-FL limits both dendritic branching and dendritic spine density in this experimental paradigm. In contrast, compared to control-transfected neurons, EphA7-T1-transfected neurons had dendrites that trended more complex (Fig.2A,B) and had significantly higher dendritic spine density (control= 0.708 ± 0.03 spines/µm control; EphA7-T1-transfected= 0.833 ± 0.04 spines/µm, p=0.020, Fig.2C,D). Thus, increasing EphA7-T1 expression had a modest effect on dendritic complexity but promoted dendritic spine formation. These results demonstrate that EphA7-FL and EphA7-T1 differentially affect dendritic branching and spine density in developing cortical neurons.

**Figure 2.**
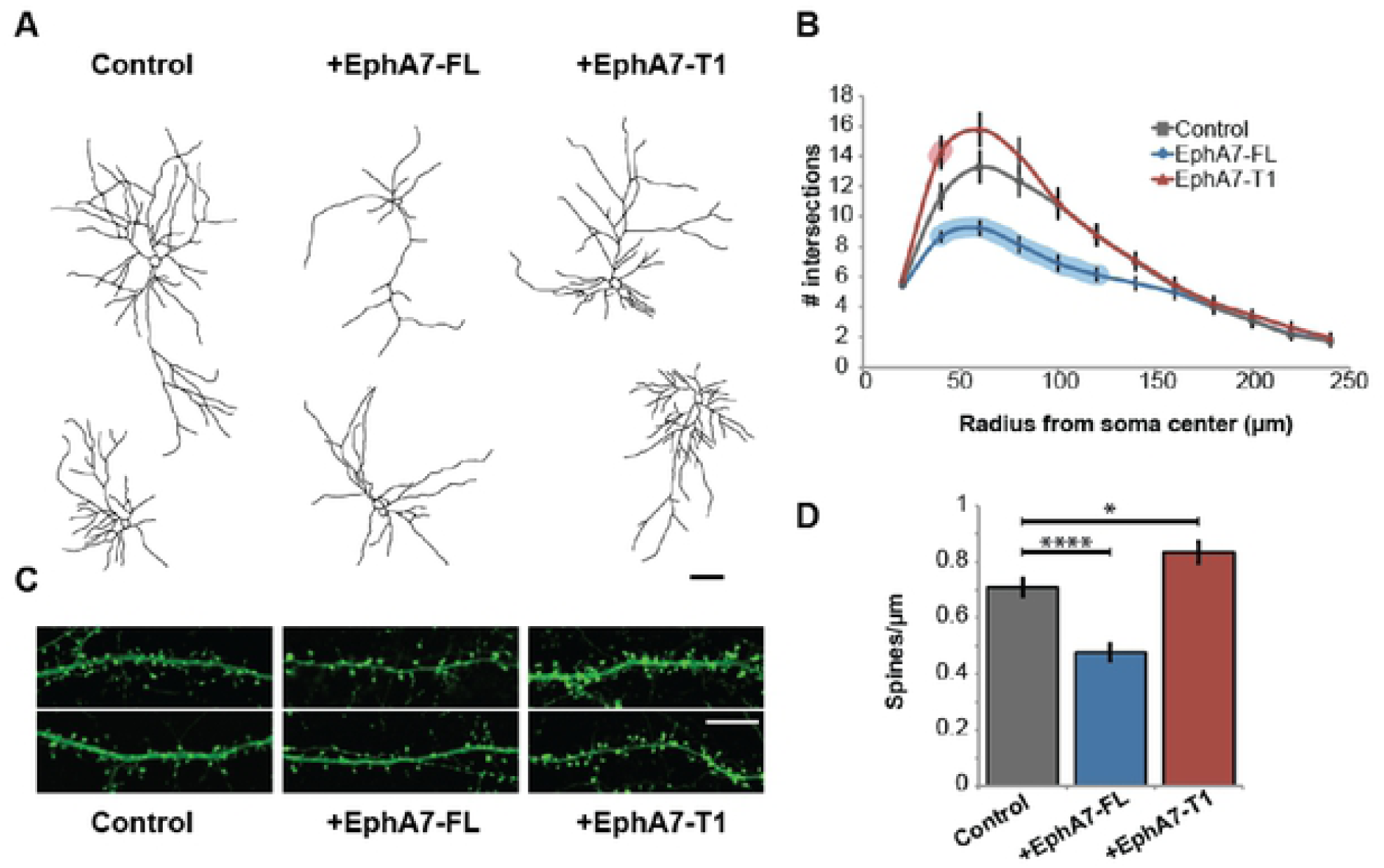
Overexpression of EphA7-FL or EphA7-T1 resulted in distinct shifts in dendritic branching and spine density. Primary cortical neurons transfected at DIV7 with actin-GFP plus control, EphA7-FL-HA, or EphA7-T1-myc plasmids were analyzed at DIV21. A. Representative traces of DIV21 control- (left), EphA7-FL-HA- (middle), or EphA7-T1- myc- (right) transfected cortical neurons (Scale bar, 50µm). B. Data from Sholl analysis of dendrites from primary cortical neurons transfected with actin-GFP plus control (gray), EphA7-FL-HA (blue), or EphA7-T1-myc (red). Compared to control, statistically significant differences in dendritic complexity, represented as number of intersections, are indicated by the shaded overlays of the same color over a line (40-120µm from soma for EphA7-FL and 40µm from soma for EphA7-T1). C. Representative images of dendrites with spines from neurons transfected with actin-GFP plus control (left), EphA7-FL-HA (middle), or EphA7-T1-myc (right) (Scale bar, 10µm) D. Compared to control-transfected neurons (gray), EphA7-FL-transfected neurons (blue) had lower dendritic and spine density EphA7-T1-transfected neurons (red) had higher dendritic (*p<0.05, ****p<0.0001.)

### Signaling downstream of EphA7 varies between elaboration of dendrites and production of spines

Our previous study suggested that dendrite repulsion induced by the EphA7 ligand, ephrin-A5, required downstream inhibition of mTOR via Tsc1, similar to a reported mechanism of EphA-guided axon retraction (Nie et al. 2010; Clifford et al. 2014). Since inhibition of mTOR is dependent on upstream phosphorylation and EphA7-FL is a kinase, we explored whether inhibition of mTOR via rapamycin could rescue dendritic phenotypes observed in *EphA7*^*-/-*^ neurons. To this end, WT and *EphA7*^*- /-*^ mice were treated with rapamycin between P5 to P22 and dendritic morphology was visualized using Golgi staining. Consistent with published findings (Clifford, et al. 2014), dendrites were more complex in vehicle-treated *EphA7*^*-/-*^ neurons compared to vehicle-treated WT neurons, (Fig.3A,B). Notably, this phenotype was rescued with rapamycin treatment, as dendrites of rapamycin-treated *EphA7*^*-/-*^ neurons were indistinguishable in complexity to vehicle-treated WT neurons (Fig.3A,B). These data indicate the *EphA7*^*-/-*^ dendritic branching phenotype arises from uninhibited mTOR activity in cortical neurons, evidenced by the phenotypic reversal following rapamycin treatment.

**Figure 3.**
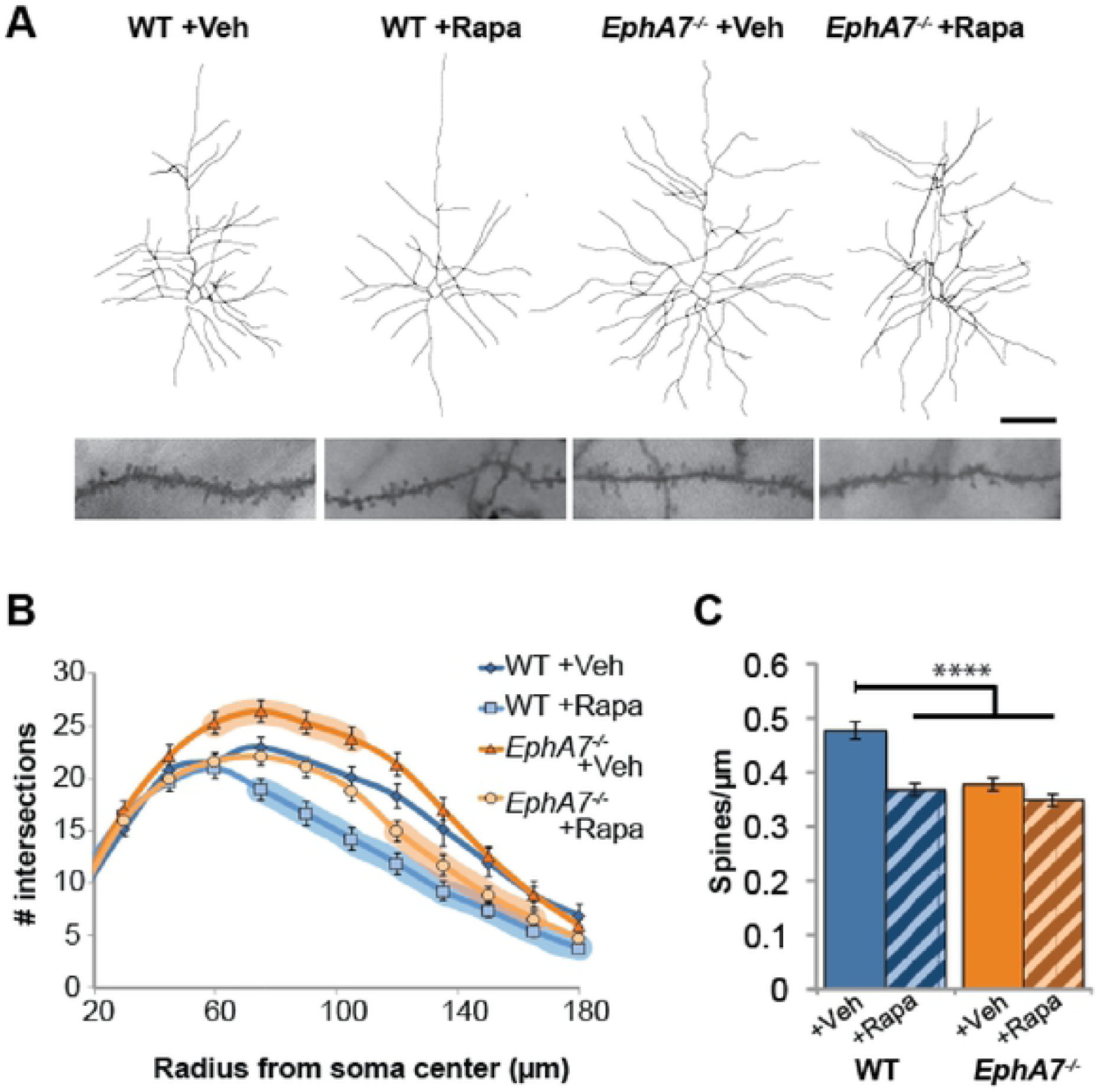
EphA7 and mTOR signaling intersect in impacting dendrite branching, but not in establishing dendritic spine density. A. Representative traces of cells (top) and images of dendrites with spines (bottom) of Golgi-stained deep layer pyramidal neurons from P22 WT (left) or *EphA1*^-/-^ (right) animals treated postnatally (from P5-P22) with vehicle (left and second from right) or rapamycin (second from left and right). (Scale bar:µm for neuronal traces;µm for dendrites). B. Data from Sholl analyses of dendrites from WT/vehicle (darker blue), WT/rapamycin (lighter blue), *EphA*7^-*/-*^/vehicle (darker orange), or *EphA*7^-/-^/ rapamycin (lighter orange) neurons. Statistically significant differences from control (WT/vehicle) are indicated by shaded overlays of the same color over a line (75- 180µm from soma for WT/rapamycin; 60-l20µm from soma for *EphA*7^-/-^/vehicle; l20- l 65µm from soma for EphA7^-/-^/ra pamy cin). C. Compared to WT/vehicle neurons (blue), both WT/rapamycin neurons (striped blue) and EphA7^-/-^/ vehicle neurons (orange) had less dense dendritic spines. There was no difference in spine density between vehicle (orange) and rapamycin (striped orange) treated *EphA7*^-/-^ neurons (For Sholl analyses: shaded overlays represent p<0.05; for spine densities ****p<0.0001.)

The potential for rapamycin to rescue *EphA7*^*-/-*^ dendritic spine phenotype was also examined. *EphA7*^*-/-*^ neurons treated with rapamycin displayed no difference in spine density compared to vehicle-treated *EphA7*^*-/-*^ neurons (*EphA7*^*-/-*^ /veh= 0.38 ± 0.01, *EphA7*^*-/-*^ /rapa= 0.35 ± 0.01, p=0.366, Fig.3A,C), demonstrating that mTOR is not implicated in EphA7-mediated dendritic spine formation. However, comparing WT neurons treated with vehicle or rapamycin, mTOR inhibition did result in decreased dendritic spine density (WT/veh= 0.48 ± 0.02, WT/rapa= 0.37 ± 0.01, p<0.0001, Fig.3A,C). Therefore, for cortical neurons, mTOR appears to be acting downstream of a signaling pathway other than EphA7 during dendritic spine formation, since rapamycin treatment affected WT dendritic spine density, but did not rescue the observed *EphA7*^*-/-*^ dendritic spine phenotype. Together, these results suggest different pathways in EphA7 downstream signaling during dendritic arborization, in which mTOR repression is implicated, versus dendritic spine formation, in which mTOR repression is not implicated.

### EphA7-FL and EphA7-T1 interact to modulate signaling and are co-expressed in cortical synaptosomes

These data support a scenario in which early in cerebral cortical development EphA7-FL acts in the canonical manner of Eph receptors, involving kinase-dependent forward signaling that inhibits mTOR and initiates cellular retraction, thus restricting dendritic arborization in cortical neurons (Davis et al. 1994; Drescher et al. 1995; S. Holland et al. 1996; Hansen et al. 2004; Soskis et al. 2012, Clifford et al. 2014). There is less clarity on how EphA7-T1, which lacks the intracellular kinase domain, acts in promoting dendritic spines. One study posited EphA7-T1 may act as a dominant negative by heterodimerizing with EphA7-FL, but an interaction was never shown (Holmberg et al. 2000). To determine whether that hypothetical interaction can actually occur, HEK cells were co-transfected with EphA7-FL-HA and EphA7-T1-myc expression vectors and co-immunoprecipitation of either isoform was performed followed by western blotting for total EphA7 and each respective epitope tag. EphA7- FL and EphA7-T1 did co-immunoprecipitate, clearly demonstrating that EphA7-FL and EphA7-T1 can directly bind to one another (Fig.4A). To address a proposed dominant negative role, manipulating the ratio of EphA7-T1 to EphA7-FL to mimic profiles in the developing cortex resulted in decreased tyrosine phosphorylation of EphA7-FL following ephrin-A5 stimulation (Fig.4B, Holmberg et al. 2000). These results demonstrate that EphA7-FL and EphA7-T1 can assemble when co-expressed in cells, and, under those circumstances, that EphA7-T1 can act to inhibit canonical EphA7 forward signaling by blocking EphA7-FL phosphorylation.

**Figure 4.**
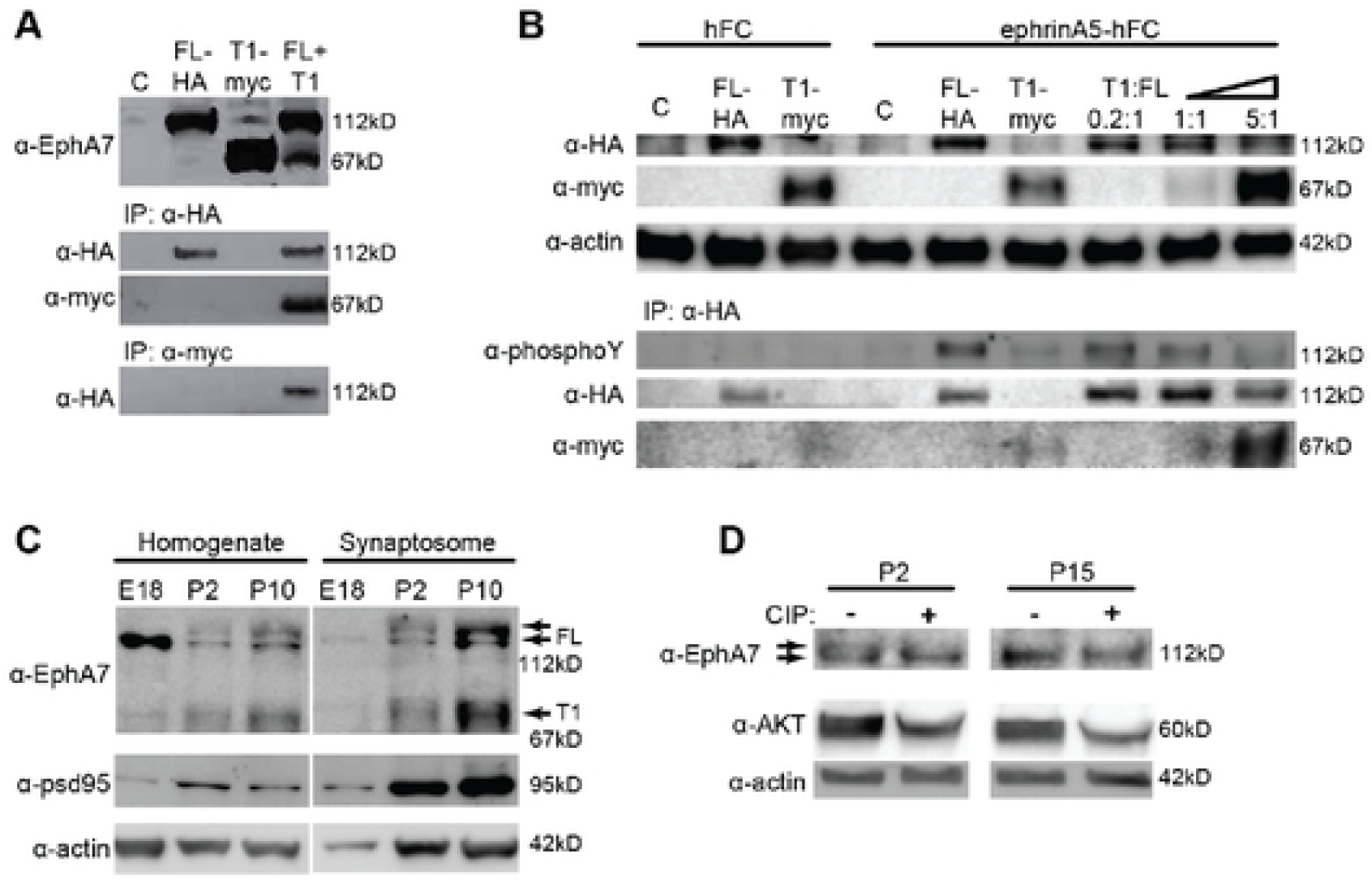
EphA7-FL and EphA7-T1 proteins interact and are colocalized at cortical synapses. A-B, HEK cells were transfected with control, EphA7-FL-HA, EphA7-T1-myc, or both EphA7-FL-HA and EphA7-T1-myc at a 1:1 ratio (A) or at increasing EphA7-T1 :FL ratios (B). A. Transfected cells were lysed and immunoprecipitated with HA or myc antibodies; western analysis detecting either total EphA7 or respective epitope tags revealed that EphA7-FL-HA and EphA7-T 1-myc protein directly interacts. B. Transfected cells were stimulated for 15 min. with clustered ephrin-AS-hFC, or hFC control. Lysates were subjected to immunoprecipitation with HA antibody to pull down FL protein, followed by detection of phospho-tyrosine, EphA7-T1-myc, and EphA7-FL-HA, which revealed that EphA7-FL tyrosine phosphorylation was decreased with increasing amounts of EphA7-T1 binding. Actin was used as loading control. C. Representative western blot of cortical homogenate and cortical synaptosomes prepared from E18, P2, or P1 Orat (all samples were run on the same gel and detected on the same membrane, lanes between homogenate and synaptosomes samples were digitally removed). EphA7 detection in cortical homogenate reveals that the intensity of EphA7-FL bands decreases with developmental time, while the intensity of EphA7-T1 and PSD95 bands increase with time (left). EphA7-FL and EphA7-T1 are detected in P2 synaptosomes, and are enriched in synaptosomes at P10, similar to PSD95 (right). Actin was used as a loading control for homogenates. D. Cortical synaptosomes from P2 (left) and P15 (right) were mock treated or treated with CIP to remove phosphate groups. A downward shift ofEphA7 EphA7-FL band at P2, but not P15, suggests that EphA7-FL phosphorylation occurs at the earlier, but not later timepoint. Akt was a positive control and actin was a loading control (all samples were run on the same gel and detected on the same membrane, lanes between P2 and P15 samples were digitally removed).

To look in developing cortex *in vivo*, protein levels of EphA7-FL and EphA7- T1 were assessed in both tissue lysates and synaptosomal fractions at E18, P2, and P10. Western blot analysis of cortical homogenates demonstrated that the intensity of EphA7-FL bands (predicted 112kD) decreased over developmental time, while the intensity of the EphA7-T1 band (predicted 67kD) increased during the same period (Fig.4C, left). In cortical synaptosomes, both EphA7-FL or EphA7-T1 were below levels of detection at E18, however both proteins were present at P2 and considerably enriched at P10. Thus, both EphA7-FL and EphA7-T1 are at cortical synapses as they mature and spines become more numerous (Fig.4C, right).

On western blots, the signal corresponding to EphA7-FL from cortex appeared broader than a single, homogeneous band, which is a result of post-translational modifications, including phosphorylation (Ciossek et al. 1995, data not shown). If EphA7-T1 acts as dominant negative to EphA7-FL, then the increase in EphA7-T1 levels in cortical synapses during development should result in less phosphorylated EphA7-FL *in vivo*. To investigate whether there are changes in EphA7-FL phosphorylation at synapses over time, synaptic fractions from P2 or P15 cortices were treated with calf intestinal phosphatase (CIP) prior to electrophoresis in order to eliminate phosphate groups. At P2, CIP-treatment caused a downward shift of the EphA7-FL band compared to the untreated sample, indicating that EphA7-FL is phosphorylated at P2 cortical synapses (Fig.4D, left). In contrast, there was no shift in the EphA7-FL band upon CIP treatment at P15 (Fig.4D, right), consistent with the idea that high levels of dominant-negative EphA7-T1 at synapses prevent phosphorylation at this stage. These results suggest that increasing levels of EphA7-T1 during development moderate EphA7-FL actions, reducing repulsive activity.

## Discussion

In summary, discrete roles for two EphA7 isoforms, EphA7-FL and EphA7- T1, are described during cortical dendrite development. Distinct expression patterns exist that correspond with developmental events: EphA7-FL is most highly expressed in the cortex just before birth, when dendrites are extending and dendritic spines are sparse whereas EphA7-T1 expression increases postnatally and peaks in the second postnatal week, a period of rapid synaptogenesis and spine formation, and remains high into adulthood, when dendrites and spines are plastic (Fig.1). Over-expression of EphA7-FL or EphA7-T1 *in vitro* mirrored *in vivo* loss-of-function phenotypes: while deletion of all *EphA7* isoforms resulted in more complex dendrites and fewer dendritic spines *in vivo*, elevated EphA7-FL expression resulted in less complex dendritic arbors and fewer dendritic spines, whereas increased EphA7-T1 caused a modest effect on branching but more dendritic spines (Fig.2, Fig.3, Clifford et al. 2014). Furthermore, EphA7-dependent dendritic elaboration is dependent on the repression of mTOR. In contrast, dendritic spine formation can be impacted by mTOR but EphA7-T1 is not acting through mTOR (Fig. 3). In addition, direct binding of EphA7-T1 to EphA7-FL occurs and influences levels of EphA7-FL phosphorylation (Fig. 4A,B). Finally, both isoforms are present at cortical synapses, collaborating to mediate signaling (Fig.4C,D).

Selective manipulation of native levels of EphA7-FL and EphA7-T1 is the clear next step and would likely provide valuable insights into isoform function at cortical synapses. Unfortunately, several attempts revealed that the EphA7 locus is largely inaccessible to CRISPR manipulation. Moreover, because EphA7-FL and EphA7-TR are identical through their entire extracellular extent and there are currently no reliable, selective protein reagents for EphA7 isoforms, transient knock-down methods could not be confidently used with current means of neuronal transfection. Thus, technical advances are required for these studies to advance.

Nonetheless, these results, when taken together, support a model in which EphA7-FL, present early in corticogenesis, restricts dendritic outgrowth through kinase-based forward signaling and repression of mTOR. Then, postnatally, EphA7- T1 levels rise, resulting in co-expression with EphA7-FL and heterodimers of the two isoforms at cortical synapses; EphA7-FL phosphorylation is inhibited and EphA7 repulsive signaling is reduced to promote formation of dendritic spines and the synapses they form with other cells (Fig.5).

**Figure 5.**
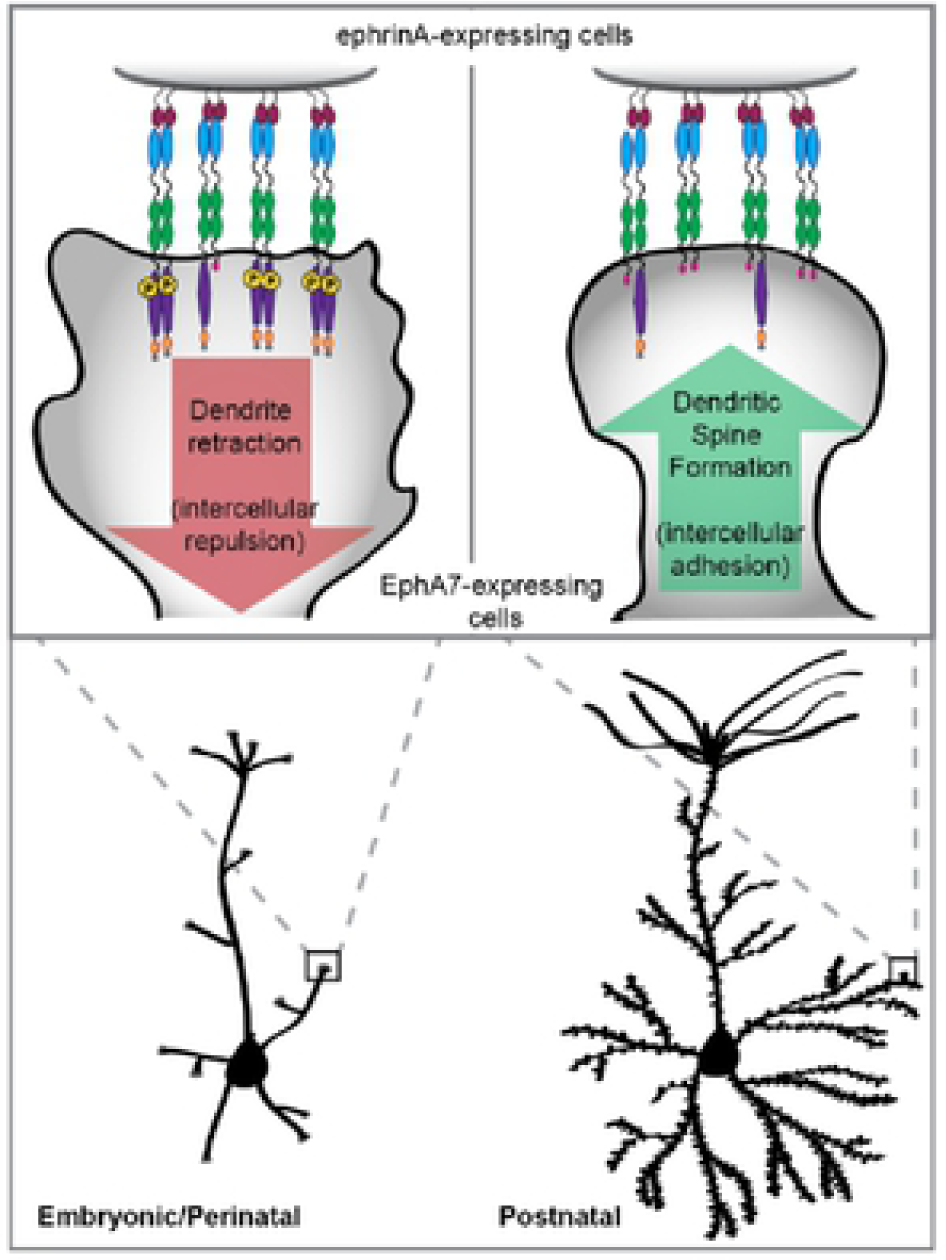
Model of EphA7 function in cortical dendrite development. Our results support a model where full-length (FL) and truncated (T1) EphA7 isoforms have distinct roles in cortical dendrite elaboration and spine formation. As neurons reach their final positions within the cortical plate, they are morphologically simple. Before and around the time of birth, dendrites extend and branch to meet incoming axons (bottom left). During this time, EphA7-FL is highly expressed. Interaction with ephrin-A-expressing cells results in EphA7-FL phosphorylation and initiates forward signaling, which consequently inhibits mTOR activity and dendritic growth (top left). In postnatal cortex, morphologically complex neurons are forming connections (bottom right). EphA7-T1 is highly expressed at this time. Therefore, interaction with ephrin-A-expressing cells no longer initiates repulsive forward signaling, but facilitates intercellular adhesion and synaptogenesis, thus promoting the formation of dendritic spines (top right).

EphA7 joins other examples of endogenous dominant negative regulation in the brain. Notably, a truncated TrkB neurotrophin receptor (TrkB-T1) inhibits full-length TrkB (TrkB-FL) activation by heterodimerization (Eide et al. 1996; Fryer et al. 1997). TrkB-T1 also has functions that are independent of TrkB-FL, including BDNF trafficking, neurite extension, and intracellular kinase regulation (reviewed in Fenner 2012). It seems likely that EphA7-T1 also has functions independent of EphA7-FL, perhaps even acting in cell adhesion.

These results highlight the importance of interpreting results with an eye toward the possibility of various receptor and ligand isoforms, including some with divergent outcomes. More broadly, several Eph receptors and ephrin ligands produce alternative mRNA transcripts and proteins of varying sizes: molecular heterogeneity that is largely neglected when assessing Eph/ephrin function (Ensembl release 89, Yates et al. 2016; NCBI Gene, Coordinators 2016). The implications of such molecular diversity warrant attention, especially as Eph receptors are increasingly discussed as potential therapeutic targets for clinical conditions.

## Materials and Methods

### Animal Husbandry

All animal use and care was in accordance with institutional, Georgetown’s GUACUC protocols #2016-1175 (mice) and #2016-1237 (rats), and federal guidelines. Control mice, either CD-1 or C57Bl/6, and rats, Sprague Dawley, were purchased from Charles River. Mice mutant for EphA7 (Rashid et al. 2005) were provided by U. Drescher (King’s College, London, UK), were backcrossed onto the C57Bl/6 strain (5-15 generations), and bred as homozygotes as previously described (Miller et al. 2006; Clifford et al. 2011). Body sizes of wild type WT and mutant animals treated with rapamycin were similar on the days of analyses. The day of the vaginal plug was considered embryonic day 0.5 (E0.5) and the day of birth, postnatal day 0 (P0). Timed pregnant females were euthanized, brains of pups were dissected and either dissociated for cell culture or lysed for protein or mRNA analysis. Postnatal animals were euthanized and brains were dissected for protein or mRNA analysis, or fixed, frozen, and sectioned, or were subjected to Golgi staining.

### Plasmids

Plasmids used in these experiments: actin-GFP, described previously (Fischer et al. 1998); pIRES2-eGFP (referred to as “empty vector” or “control”, Clontech); EphA7-FL-HA (“EphA7-FL-HA”); EphA7-T1-myc (“EphA7-T1-myc”). EphA7-FL-HA and EphA7-T1- myc expression plasmids were created by subcloning cDNA corresponding to either isoform into pIRES2-EGFP (Clontech) and inserting epitope tags via polymerase chain reaction with Pfu DNA polymerase (Promega). Primers for HA insertion into pIRES2- EGFP-FL: forward: 5’-ACCAGACTATGCCCCTGACTTCACTGCCTTCTGTTC-3’, reverse: 5’-ACATCATAGGGATAAGTGCTCTGGTCCAGAAGGAAGC-3’. Primers for myc insertion into pIRES2-EGFP-T1: forward: 5’- ATCTGAAGAGGACTTGTAAACCGCAACAATAACTGTTTAAGAG-3’, reverse: 5’- ATCTGCTTTTGCTCTAAAACTGACAGGTGCTCATTTGTTAC-3’.

### Golgi Staining and Analyses

Golgi Staining was performed according to manufacturer instructions (FD Neurotechnologies). Briefly, brains from P22 mice were dissected, weighed, rinsed, and incubated in solution A/B for two weeks. Following a week-long clearing step in solution C, brains were frozen in a dry ice/isopentane bath, and sectioned on a cryostat at 120- 180μm. Sections were mounted on gelatin-coated slides and dried overnight before being developed in D/E solution, dehydrated, and mounted using Permount (Sigma).

#### Analysis of dendritic extent in vivo

The NeuronJ plug-in (Meijering et al. 2004) for ImageJ (NIH) was used to trace spiny pyramidal neurons in deep layers of mouse motor cortex for examination of dendritic arborization. Two or three neurons were traced from each animal, capturing the entire basal dendritic tree and the first 200 μm of the apical dendrite with oblique branches. Sholl analysis quantified dendritic complexity (Sholl 1953). Sholl analysis: Concentric circles of increasing radius (15-20μm intervals) were centered on the soma and numbers of dendritic intersections were counted along each circle at a given radius. The average and standard error of the mean (SEM) are reported, referring to the total number of neurons measured from a given group (specified in figure legends). Groups were compared to WT first using two-way (WT±rapa vs *EphA7*^*-/-*^ ±rapa) ANOVA, followed by Fisher’s LSD post-hoc analysis in Prism 6 (GraphPad).

#### Analysis of dendritic spine density in vivo

Spiny pyramidal neurons in deep layers of mouse motor cortex were examined. For each cell, a 50 μm segment of a secondary basal or apical oblique branch was identified, the number of dendritic spines within each segment was recorded, and density was calculated as number of dendritic spines divided by 50 μm. Average and SEM are reported, referring to the total number of cells analyzed within a given group. Comparisons were made using two-way (WT/veh vs WT/rapa vs *EphA7*^*-/-*^ /veh vs *EphA7*^*-/-*^ /rapa) ANOVA, followed by post-hoc analysis with Dunnet correction (GraphPad Prism 6).

### In Vivo *Rapamycin Treatment*

Before each injection, animals were weighed and treatment doses were prepared (6 mg/kg) by diluting in sterile vehicle (2.5mL 5% polyethylene glycol, 2.5mL 5% Tween-80, and 45mL sterile, deoionized water, filtered with Steriflip (Millipore)). WT or *EphA7*^*-/-*^ pups were treated with rapamycin (LC labs) via intraperitoneal injection 3 days per week (MWF) from P5-6 until P22-23, when they were euthanized. Upon euthanization, animals were weighed, and brains were dissected and subjected to Golgi staining. Consistent with other reports in young mice, rapamycin-treated animals weighed less than vehicle-treated animals, with no significant differences in brain weight (Anderl et al. 2011). There were no differences in weight due to genotype during the course of the experiment (Supp. Fig. 3).

### Primary Cell Culture

Primary cortical neurons were prepared from E18.5 rat embryos. One day before culture, acid-washed, glass coverslips were coated with 100 μg /mL poly-d-lysine in sterile PBS and incubated overnight at room temperature. The next day, coverslips were washed with sterile water prior to plating. An E18.5 pregnant rat was euthanized with CO_2_ asphyxiation and pups were collected. Dorsal cortex was immediately dissected in 1x HBSS+ (Hank’s Balanced Salt Solution plus 10mM HEPES and 1% penicillin/streptomycin). Chopped tissue was washed in fresh HBSS^+^, then incubated in 0.125% trypsin/HBSS+ for fifteen minutes at 37° Celsius. Tissue was rinsed three times with HBSS and triturated in neuronal growth media (Neurobasal supplemented with 2% B27, 1% Glutamax, and 1% penicillin/streptomycin). Cells were plated at a density of 150,000 cells per well of a 12- well dish and maintained at 37° C, 5% CO_2_ in a humidified environment. Cultures were fed by replacing half of the media volume with fresh growth media twice per week.

### Cell Line Culture

HEK293 cells were grown in DMEM (Dulbecco’s Modified Eagle Medium) supplemented with 10% fetal bovine serum and 1% penicillin/streptomycin, and maintained at 37° C with 5% CO_2_ in a humidified environment. Cells were passaged at no more than 80% confluency by brief trypsinization at room temperature, followed by serum inactivation and plating in fresh growth media.

### Quantitative RT-PCR

Total RNA was isolated from cerebral cortical tissue of E12.5, E14.5, E16.5, E18.5, P0, P5, P10, P21 and P60 mice or from cultures of mouse primary cortical neurons at days *in vitro*(DIV) 1, 2, 3, 5, 8, 10, and 13 using the Tri-Regent Kit (Sigma-Aldrich). cDNA was synthesized using the First-Strand cDNA Synthesis Kit (Invitrogen). Primers specific for mouse EphA7-FL, EphA7-T1, and U6 (EphA7-FL forward: 5′- CGTCTGAAGATGCTGGTGAAAGCA-3′, EphA7-FL reverse: 5′- ACTGAAGCTCTTGATGTGCTCGGA-3′; EphA7-T1 forward: 5′- TAAGGACCTGGATTCCTTCAGCGA-3′, EphA7-T1 reverse: 5′- CTGCATGCTTCTGGGTGCATTGAA-3′; U6 forward: 5′-CTCGCTTCGGCAGCACA- 3′, U6 reverse: 5′-AACGCTTCACGAATTTGCGT-3′) were designed and used in quantitative real-time RT-PCR (qRT-PCR). qRT-PCR was performed using the 2× SYBR Green PCR Master Mix (Bioline) with the CFX96 detection system and analyzed with CFX Manager softward (Bio-rad). Transcript levels were normalized to U6 and converted to percent maximum relative expression for EphA7-FL and EphA7-T1 separately. Each individual experiment was performed with experimental triplicates and each timepoint was analyzed from three separate animals. Reported values are the average and SEM of 3 experiments (animals).

### Transfection

#### HEK cell transfection

HEK cells were transfected at 50-60% confluent in complete growth medium, using X-tremeGENE (Roche), according to manufacturer’s instructions. For one well of a 6-well dish, 2 μg total DNA was transfected at 1:1 DNA:reagent ratio, where 0.5 μg /well of each expression plasmid was added plus empty vector to total 2 μg. Transfection cocktail was added directly to the well and left until cell lysis.

#### Neuron transfection

Primary cortical neurons were transfected at DIV7 with Lipofectamine 2000 (ThermoFisher). One day before transfection, conditioned media was prepared by removing half the volume of media per well and mixing with equal volume fresh neuronal growth media, then supplementing the neurons with fresh media. On the day of transfection, transfection cocktails were prepared according to manufacturer instructions, by mixing (per well of 12-well dish) 200uL non-supplemented Neurobasal media, 0.1 μg /kB of desired expression plasmid(s), and 1uL transfection reagent. Transfection cocktails were added to neurons and allowed to rest at 37° C Celsius for 60- 90 minutes. Transfection media was completely removed and replaced with conditioned media, and neurons were returned to 37° C.

### Ligand Binding Assay

To determine whether EphA7 expression constructs bind to ephrin ligand, HEK cells were transfected with empty vector, EphA7-FL-HA, or EphA7-T1-myc. After 24hrs, cells were fixed in 4% paraformaldehyde for 20 minutes, then rocked at room temperature for at least one hour, while incubating with ephrinA5 conjugated to human FC (R&D) at a concentration of 3mg/ml in 0.5x blocking buffer (0.01% Triton X-100, 1% BSA) in PBS. Cells were fixed again and subjected to immunocytochemistry (Davis et al. 1994)

### Fluorescent Immunocytochemistry

The following primary antibodies were used: rabbit anti-GFP (ThermoFisher, 1:3000 dilution); rabbit anti-NeuN (CST, 1:1000); rabbit anti-GFAP (CST, 1:1000); mouse anti- PSD95 (Sigma, 1:1000); rabbit anti-HA (CST, 1:1000); rabbit anti-myc (CST, 1:1000); goat anti-EphA7 (R&D, 1:500); goat anti-human (ThermoFisher, 1:1000). Hoechst staining (ThermoFisher, 1:15,000) was also performed. Appropriate Alexafluor-conjugated secondary antibodies (ThermoFisher) were used at a dilution of 1:500. For live imaging of transfected HEK cells, goat anti-EphA7 (R&D) was diluted 1:500 in pre-warmed, serum free DMEM and growth media was replaced with anti-EphA7/media cocktail for at least two hours at 37°C before fixation and application of secondary antibody. After their respective treatments, cells were fixed for 20 min with 4% paraformaldehyde/4% sucrose at room temperature. Cells were permeabilized in 0.1% Triton X-100 in PBS, before blocking in 0.1% Tx-100 and 10% BSA in PBS. Cells were incubated with primary antibody diluted in 0.1% Tx-100 and 1% BSA in PBS overnight at 4°C, then with secondary antibody for at least 1 hour at room temperature before brief application of Hoechst stain in PBS, followed by rinsing and mounting with Fluoromount-G (SouthernBiotech).

### *In vitro* analysis of dendritic extent and dendritic spine density

Neuron morphology was visualized by ICC for GFP. Neurons were selected for analysis if they were spiny, and displayed 4-7 primary dendrites, with a prominent “apical” dendrite. For each experiment (performed from three separate culture preparations), at least twenty neurons were traced from two coverslips for analysis of dendritic morphology and ten neurons were analyzed from each condition for dendritic spine counts.

#### Analysis of dendritic extent

Sholl analysis was performed as described above. The average and standard error of the mean (SEM) are reported for one representative experiment, referring to the total number of neurons measured in that experiment (specified in figure legends). Groups were compared to control-transfected neurons first by one-way ANOVA, followed by Fisher’s LSD post-hoc analysis in Prism 6 (GraphPad).

#### Analysis of dendritic spine density

Dendritic spine density was calculated as described above. Average and SEM are reported for one representative experiment, referring to the total number of neurons measured in that experiment. Comparisons were made using one-way ANOVA, followed by Dunnet adjusted post-hoc analysis (GraphPad Prism 6).

### RNAscope multiplex fluorescent in situ *hybridization assay*

RNAscope probes were designed to detect mouse EphA7 mRNA for either full length (FL, Mm-Epha7-tv1tv3, targeting sequence: 2189-3239 of NM_010141.4) or truncated isoform (T1, Mm-Epha7-tv2, targeting sequence: 2064-3253 of NM_001122889.1). RNAscope Multiplex Fluorescent Assay on cultured cells and fixed frozen tissue sections was performed according to manufacturer’s instructions (Advanced Cell Diagnostics, ACD, Hayward, CA). Dissociated cortical neurons were cultured on glass cover slips and fixed in 4% formaldehyde solution for 30 min (room temperature) at DIV 7, 14, and 21. Fixed cells were dehydrated and stored in 100% ethanol at -20 °C for up to 6 months. Prior to hybridization, cells were rehydrated, permeabilized with 1x PBS containing 0.1 % Tween-20 for 20 min and treated with protease III (1:10, ACD) for 20 min. For RNAscope fluorescent assay on fixed frozen tissue, the slides containing 12.5 μm -thick coronal sections of mouse brains, collected at P0 and P10 were submerged into boiling 1x target retrieval buffer (ACD) for 5-10 min, followed by protease III treatment (ACD) for 30 min at 40 °C. Following ACD protocol for hybridization and signal amplification steps, the cover slips with cultured cortical cells or slides with brain tissue were either mounted with Fluoromount-G (SouthernBiotech) or subjected to a subsequent immunocytochemistry. Images were acquired with the same parameter settings using Zeiss LSM 880 confocal microscope. mRNA puncta were quantified using Imaris software (Bitplane, Oxford Instruments). For cortical culture, 20 neurons from 3 separate experiments were analyzed for each time point in vitro. For in vivo experiments, mRNA puncta were quantified in sections containing motor cortex, 2-3 sections per animal, 3-4 animals per age group. Comparisons between EphA7-FL and EphA7-T1 in all cases were made using two-way ANOVA, followed by Holm-Sidak (in vivo) or Dunnet (in vitro) corrected post-hoc analysis. Comparison of FL:T1 ratio per cell was made using one-way ANOVA, followed by Dunnet corrected post-hoc analysis.

### Co-immunoprecipitation Assay and ephrin-A5 Treatment in HEK cells

For co-immunoprecipitation, protein A/G agarose beads (Santa Cruz) were washed in immunoprecipitation buffer (1% NP-40, 50 mM Tris-HCl, pH 8.0, 150 mM NaCl) and incubated with primary antibody at 1:200 dilution (rabbit anti-HA or anti-myc, CST) for ≥1hr, then rinsed in IP buffer. For ephrin-A5 treatment (Fig. 4B), ephrin-A5 conjugated to human FC (ephrinA5-hFC, R&D) or hFC control (R&D) was pre-clustered with goat anti-human antibody (Abcam) for one hour at 37°C before being added to transfected HEK cells (5 μg /ml) for 15 minutes. Treated or untreated transfected HEK cells were lysed in cold IP buffer with protease inhibitors (Roche), and centrifuged for 10 minutes and 5,000 rpm at 4°C. Supernatant was collected and measured via Bradford Protein Assay (Bio-rad). 200 μg total protein was incubated with prepared agarose beads in IP buffer overnight at 4°C. Samples were washed in cold IP buffer before being eluted in 2X Laemmli sample buffer with DTT, boiled, and separated via SDS-PAGE.

### Synaptosomal Fractionation, and Phophatase Tx of Cortical Lysate

Crude synaptosomal fractions were collected from embryonic or postnatal rat cortex using the Syn-PER Extraction Reagent (ThermoSci) according to manufacturer’s directions. Briefly, dissected cortical tissue was weighed and gently homogenized in ice cold Syn-PER reagent plus protease inhibitors (10mL/gram of tissue). Samples were centrifuged at 1200 x *g* for 10 minutes at 4°C and supernatants were collected (homogenate sample). For synaptosomes, samples were centrifuged again at 15,000 x *g* for 20 minutes at 4°C and the pellet was resuspended in Syn-PER reagent (synaptosome sample) or the supernatant was collected (cytosolic sample). For phosphatase treatment, synaptosomal and cytosolic factions were treated with CIP (0.5units/μg, NEB) in CutSmart buffer (NEB) for 60 minutes at 37°C. Samples were mixed with 2X Laemmli sample buffer with DTT, boiled, and separated via SDS-PAGE.

### Western Blotting

Cell lysates were collected as described and denatured by boiling in Laemmli sample buffer with DTT before SDS-PAGE separation on 12% pre-cast gels (Bio-rad). Western blots were performed using the Bio-Rad Mini TransBlot and TransBlot-Turbo Transfer System, with the Bio-Rad PVDF Transfer Kit. Chemiluminescent imaging was performed on the ImageQuant LAS4000 biomolecular imager (GE Healthcare). The following primary antibodies were used: rabbit anti-EphA7 (Santa Cruz, 1:500), mouse anti-HA (Sigma, 1:1000), rabbit anti-HA (CST, 1:1000), mouse anti-myc (Sigma, 1:1000), rabbit anti-myc (CST, 1:1000), mouse anti-actin (Sigma, 1:2500), rabbit anti-phospho-tyrosine (CST, 1:1000), mouse anti-psd95 (CST, 1:1000), rabbit anti-Akt(pan) (CST, 1:1000). Appropriate HRP-conjugated secondary antibodies (Jackson ImmunoResearch) were used at a dilution of 1:2500.

## Acknowledgments

We thank J. Mejia for technical assistance and animal husbandry. We thank L. Orefice for helpful scientific advice and discussion. We thank T. Cung and B. Burgess for assistance with data collection and analyses.

## References

Amegandjin, C.A. et al., 2016. Regional expression and ultrastructural localization of EphA7 in the hippocampus and cerebellum of adult rat. Journal of Comparative Neurology, 524(12), pp. 2462–2478.

Anderl, S. et al., 2011. Therapeutic value of prenatal rapamycin treatment in a mouse brain model of tuberous sclerosis complex. Human Molecular Genetics, 20(23), pp.4597–4604.

Angevine, J.B.J. & Sidman, R.L., 1961. Autoradiographic study of cell migration during histogenesis of cerebral cortex in the mouse. Nature, 192, pp.766–768.

Anton, E.S., Cameron, R.S. & Rakic, P., 1996. Role of Neuron-Glial Junctional Domain Proteins in the Maintenance and Termination of Neuronal Migration across the Embryonic Cerebral Wall. The Journal of Neuroscience, 16(7), pp.2283–2293.

Arlotta, P. et al., 2005. Neuronal subtype-specific genes that control corticospinal motor neuron development in vivo. Neuron, 45(2), pp.207–221.

Calabrese, B., Wilson, M.S. & Halpain, S., 2006. Development and Regulation of Dendritic Spine Synapses. Physiology, 21, pp.38–47.

Chen, Y., Fu, A.K.Y. & Ip, N.Y., 2008. Bidirectional signaling of ErbB and Eph receptors at synapses. Neuron glia biology, 4(3), pp.211–21.

Ciossek, T. et al., 1999. Segregation of the receptor EphA7 from its tyrosine kinasenegative isoform on neurons in adult mouse brain. Brain research. Molecular brain research, 74(1-2), pp. 231–6.

Ciossek, T., Millauer, B. & Ullrich, A., 1995. Identification of alternatively spliced mRNAs encoding variants of MDK1, a novel receptor tyrosine kinase expressed in the murine nervous system. Oncogene, 10, pp.97–108.

Clifford, M. a et al., 2011. EphA4 expression promotes network activity and spine maturation in cortical neuronal cultures. Neural development, 6(1), p.21.

Clifford, M. a. et al., 2014. EphA7 signaling guides cortical dendritic development and spine maturation. PNAS, pp. 3–8.

Coordinators, N.R., 2016. Database resources of the National Center for Biotechnology Information. Nucleic Acids Research, 44(Database issue), pp.D7–D19.

Davis, S. et al., 1994. Ligands for EPH-related receptor tyrosine kinases that require membrane attachment or clustering for activity. Science (New York, N.Y.), 266(October), pp.816–819.

Drescher, U. et al., 1995. In vitro guidance of retinal ganglion cell axons by RAGS, a 25 kDa tectal protein related to ligands for Eph receptor tyrosine kinases. Cell, 82(3), pp.359–70.

Eide, F.F. et al., 1996. Naturally occurring truncated trkB receptors have dominant inhibitory effects on brain-derived neurotrophic factor signaling. The Journal of neuroscience : the official journal of the Society for Neuroscience, 16(10), pp.3123–9.

Fenner, B.M., 2012. Truncated TrkB: Beyond a dominant negative receptor. Cytokine and Growth Factor Reviews, 23, pp.15–24.

Fischer, M. et al., 1998. Rapid actin-based plasticity in dendritic spines. Neuron, 20(5), pp.847–54.

Fryer, R.H., Kaplan, D.R. & Kromer, L.F., 1997. Truncated trkB receptors on nonneuronal cells inhibit BDNF-induced neurite outgrowth in vitro. Experimental neurology, 148(2), pp.616–627.

Gale, N.W. et al., 1996. Eph Receptors and Ligands Comprise Two Major Specificity Subclasses and Are Reciprocally Compartmentalized during Embryogenesis. Neuron, 17, pp.9–19.

Glausier, J.R. & Lewis, D. a, 2012. Dendritic spine pathology in schizophrenia. Neuroscience, p.http://dx.doi.org/10.1016/n.neuroscience.2012.04.0.

Guan, K.-L. & Rao, Y., 2003. Signaling mechanisms mediating neuronal responses to guidance cues. Nature Reviews Neurosci, 4, pp.941–956.

Hansen, M.J., Dallal, G.E. & Flanagan, J.G., 2004. Retinal axon response to ephrin-As shows a graded, concentration-dependent transition from growth promotion to inhibition. Neuron, 42(5), pp.717–730.

Häusser, M., Spruston, N. & Stuart, G.J., 2000. Diversity and Dynamics of Dendritic Signaling. Science, 290, pp.739–744.

Himanen, J.-P. et al., 2004. Repelling class discrimination: ephrin-A5 binds to and activates EphB2 receptor signaling. Nature Neuroscience, 7(5), pp.501–509.

Holland, S. et al., 1996. Bidirectional signalling through the EPH-family receptor Nuk and its transmembrane ligands. Nature, 383, pp.722–725.

Holland, S.J. et al., 1996. Bidirectional signalling through the EPH-family receptor Nuk and its transmembrane ligands. Nature, 383, pp.722–5.

Holmberg, J., Clarke, D.L. & Frisen, J., 2000. Regulation of repulsion versus adhesion by different splice forms of an Eph receptor. Nature, 408, pp.203–206.

Kaufmann, W.E. & Moser, H.W., 2000. Dendritic anomalies in disorders associated with mental retardation. Cerebral cortex (New York, N.Y. : 1991), 10, pp.981–91.

Koester, S.E. & O’Leary, D.D.M., 1992. Functional Classes of Cortical Projection Neurons Develop Dendritic Distinctions by Class-specific Sculpting of an Early Common Pattern. J Neuro, 12(4), pp.1382–393.

Kwiatkowski, D.J. et al., 2002. A mouse model of TSC1 reveals sex-dependent lethality from liver hemangiomas, and up-regulation of p70S6 kinase activity in Tsc1 null cells. Human molecular genetics, 11(5), pp.525–34.

Ledda, F. & Paratcha, G., 2017. Mechanisms regulating dendritic arbor patterning. Cell. Mol. Life. Sci.

Lehigh, K.M. et al., 2013. Parcellation of the thalamus into distinct nuclei reflects EphA expression and function. Gene expression patterns : GEP, 13(8), pp.454–63.

Martin, K.C. & Zukin, R.S., 2006. RNA Trafficking and Local Protein Synthesis in Dendrites: An Overview. The Journal of Neuroscience, 26(27), p.7131 LP –7134.

McConnell, S.K., 1995. Constructing the Cerebral Cortex: Neurogenesis and Fate Determination. Neuron, 15, pp.761–768.

Meijering, E. et al., 2004. Design and validation of a tool for neurite tracing and analysis in fluorescence microscopy images. Cytometry Part A, 58(2), pp.167–76.

Meikle, L. et al., 2007. A mouse model of tuberous sclerosis: neuronal loss of Tsc1 causes dysplastic and ectopic neurons, reduced myelination, seizure activity, and limited survival. The Journal of neuroscience : the official journal of the Society for Neuroscience, 27(21), pp.5546–5558.

Miller, K., Kolk, S.M. & Donoghue, M.J., 2006. EphA7-ephrin-A5 Signaling in Mouse Somatosensory Cortex : Developmental Restriction of Molecular Domains and Postnatal Maintenance of Functional Compartments. Journal of Comparative Neurology, 496, pp.627–642.

Miller, M., 1981. Maturation of rat visual cortex. I. A quantitative study of Golgiimpregnated pyramidal neurons. Journal of neurocytology, 10(5), pp.859–78.

Molyneaux, B.J. et al., 2007. Neuronal subtype specification in the cerebral cortex. Nature reviews. Neuroscience, 8(6), pp.427–437.

Mori, T. et al., 1995. Localization of novel receptor tyrosine kinase genes of the eph family, MDK1 and its splicing variant, in the developing mouse nervous system. Molecular Brain Research, 34, pp.154–160.

Nie, D. et al., 2010. Tsc2-Rheb signaling regulates EphA-mediated axon guidance. Nature neuroscience, 13(2), pp.163–72.

Noctor, S.C. et al., 2004. Cortical neurons arise in symmetric and asymmetric division zones and migrate through specific phases. Nat Neurosci, 7(2), pp.136–144.

Noren, N.K. & Pasquale, E.B., 2004. Eph receptor-ephrin bidirectional signals that target Ras and Rho proteins. Cellular signalling, 16(6), pp.655–666.

Phillips, M. & Pozzo-Miller, L., 2015. Dendritic spine dysgenesis in Autism Related Disorders. Neurosci Lett, 601, pp.30–40.

Polleux, F. et al., 1998. Patterning of cortical efferent projections by semaphorin-neuropilin interactions. Science, 282, pp.1904–6.

Polleux, F., Morrow, T. & Ghosh, A., 2000. Semaphorin 3A is a chemoattractant for cortical apical dendrites. Nature, 404, pp.567–73.

Rakic, P., 1974. Neurons in Rhesus Monkey Visual Cortex: Systematic Relation between Time of Origin and Eventual Disposition. Science, 183(4123).

Rashid, T. et al., 2005. Opposing Gradients of Ephrin-As and EphA7 in the Superior Colliculus Are Essential for Topographic Mapping in the Mammalian Visual System. Neuron, 47(1), pp.57–69.

Sala, C., Cambianica, I. & Rossi, F., 2008. Molecular mechanisms of dendritic spine development and maintenance. Acta neurobiologiae experimentalis, 68(2), pp.289–304.

Sholl, D.A., 1953. Dendritic organization in the neurons of the visual and motor cortices of the cat. Journal of Anatomy, 87(4), pp.387–406.

Sialana, F.J. et al., 2016. Mass spectrometric analysis of synaptosomal membrane preparations for the determination of brain receptors, transporters and channels. Proteomics, 16, pp.2911–2920.

Sidman, R.L. & Rakic, P., 1982. Development of the human central nervous system. In D. Adams, ed. Histology and histopathology of the nervous system. Springfield, Illinois.: C.C. Thomas, pp. 3–145.

Soskis, M.J. et al., 2012. A chemical genetic approach reveals distinct EphB signaling mechanisms during brain development. Nature neuroscience, 15(12), pp.1645–54.

Torii, M. & Levitt, P., 2005. Dissociation of corticothalamic and thalamocortical axon targeting by an EphA7-mediated mechanism. Neuron, 48(4), pp.563–575.

Tsai, L.-H. & Gleeson, J.G., 2005. Nucleokinesis in Neuronal Migration. Neuron, 46, pp.383–388.

Valenzuela, D.M. et al., 1995. Identification of full-length and truncated forms of Ehk-3, a novel member of the Eph receptor tyrosine kinase family. Oncogene, 10(8), pp.1573–80.

Vercelli, A., Assal, F. & Innocenti, G.M., 1992. Emergence of callosally projecting neurons with stellate morphology in the visual cortex of the kitten. Experimental brain research, 90(2), pp.346–58.

Whitford, K.L. et al., 2002. Molecular control of cortical dendrite development. Annual review of neuroscience, 25, pp.127–49.

Yates, A. et al., 2016. Ensembl 2016. Nucleic Acids Research, 44(D1), pp.D710–D716.

Yun, M.E. et al., 2003. EphA family gene expression in the developing mouse neocortex: regional patterns reveal intrinsic programs and extrinsic influence. The Journal of comparative neurology, 456(3), pp.203–16.

Zhu, Y. et al., 2001. Ablation of NF1 function in neurons induces abnormal development of cerebral cortex and reactive gliosis in the brain. Genes and Development, 15(7), pp.859–876.

